# Evolution of resistance to KRAS^G12C^ inhibitor in a non-small cell lung cancer responder

**DOI:** 10.1101/2023.12.18.572090

**Authors:** Jia-Hui Xu, Shi-Jia Wang, Ziming Wang, Jumin Huang, Chun Xie, Yabing Cao, Ming Chen, Elaine Lai-Han Leung

## Abstract

Despite initial therapeutic successes, most patients with non-small cell lung cancer (NSCLC) who carry the KRAS^G12C^ mutation ultimately exhibit resistance to targeted treatments. To improve our comprehension of how acquired resistance develops, we present an unprecedented longitudinal case study profiling the transcriptome of peripheral blood mononuclear cells (PBMCs) over 5 months from an NSCLC patient with the KRAS^G12C^ mutation and initial response to sotorasib followed by resistance and death. Single-cell RNA sequencing analysis uncovered notable fluctuations in immune cell populations throughout treatment with sotorasib. Specifically, we observed a decline in circulating CD8^+^CD161^hi^ T cells correlating with periods of therapeutic response, followed by a resurgence during phases of nonresponse. This study established a high-resolution atlas detailing the evolutionary trajectory of resistance to sotorasib and characterizes a CD8^+^CD161^hi^ T cells population in KRAS^G12C^ mutation patient.

## Main

Annually, China reports over four million new cancer cases, with lung cancer comprising 24.6% and being the deadliest in terms of mortality^1^. Lung adenocarcinoma (LAC), which constitutes approximately 30-35% of all lung cancer cases, is characterized by a markedly high incidence of KRAS mutations, with nearly 100% of cases exhibiting such genetic alterations^2^. Despite KRAS’s reputation as “undruggable,” the Food and Drug Administration (FDA) has approved two KRAS^G12C^ inhibitors, sotorasib and adagrasib, for advanced NSCLC with the KRAS^G12C^ mutation^3,4^. However, the CodeBreaK 200 phase III trial’s progression-free survival (PFS) for sotolacib, the first FDA-approved KRAS^G12C^ inhibitor, was less than anticipated^5^. Current investigations have elucidated multiple resistance mechanisms to KRAS^G12C^ inhibitors, including secondary mutations, reactivation of the bypass signalling pathway, acquired KRAS alterations, and the epithelial–mesenchymal transition^6-9^. Other potential mechanisms, such as other epigenetic mechanisms, gut microbiota, and immune destruction factors, remain to be investigated^10-12^.

Single-cell technologies have completely changed how we investigate the progression of tumours and medication resistance^13^, enabling the establishment of detailed immune profiles in cancers^14,15^. Longitudinal tumour sampling facilitates investigation of temporal response dynamics, offering insights into tumour heterogeneity, evolution, and development of acquired resistance^16^. Recently, our longitudinal study on Chinese NSCLC patients receiving anti-PD1 therapy, with over 30 months of follow-up, revealed that specific CD8^+^ subpopulations correlate significantly with anti-PD-1 therapy^17^. This finding inspired us to develop a robust strategy for utilizing noninvasive biopsies, and it is necessary to apply novel highdimensional and single-cell technologies to explore the reason for sotorasib resistance. Sotorasib is currently first being launched into the Chinese market in the Macau special administrative region, and this is the first clinical case report of sotorasib resistance in China. In this work, we used single-cell RNA-sequencing (scRNA-seq) technology to characterize the cellular and molecular dynamics of immune cells in one patient with NSCLC treated with sotorasib by following longitudinal sampling over a period of 5 months to gain insights into the changes in the immune cell population and their gene expression profiles over time. We sought to uncover the mechanisms behind resistance and sensitivity to targeted inhibitor therapies in NSCLC, identify immune cell subtypes that react to sotorasib, and address its treatment limitations.

To elucidate development of resistance to sotorasib, we assembled a distinctive dataset derived from 4 peripheral blood samples collected longitudinally over 5 months; to date, this is the longest follow-up time of patients who receive sotorasib treatment in China. This dataset chronicles a patient’s transition from a favourable initial response to sotorasib to disease progression and death (Fig. 1A). We conducted single-cell RNA sequencing analysis of the cells from these samples. After quality filtering, we obtained single-cell transcriptome data for 30,000 high-quality immune cells, which we classified into 41 distinct clusters (Clusters 0-40) to maximize differentiation of cell populations (Fig. 1B, Supplementary Fig. S1A, B). Following this stratification, we employed SingleR^18^ for cluster annotation, ultimately identifying 7 major cell types, including T cells (marked by *CD3D, CD3E, CD4* and *CD8A*), natural killer (NK) cells (marked by *NKG7*), B cells (marked by *MS4A1*), monocytes (marked by *CD14*), dendritic cells (DCs) (marked by *FCER1A*), macrophages (marked by *FCGR3A*) and megakaryocytes (marked by *PPBP*) (Fig. 1C). Through marker gene analysis for cell type identification, we detected the presence of megakaryocytes within our samples. Interestingly, the proportions of T cells, NK cells, B cells, and dendritic cells (DCs) increased during the drug response phase and decreased when there was no response (Figure 1D), with declines in T cells and NK cells being the most pronounced (Supplementary Fig. S1C). Monocytes increased in both response and nonresponse cycles. Subsequently, we employed the CellChat^19^ tool to dissect the intricacies of cell-to-cell communication, and discovered that complex contact among these 7 cell types, especially T cells, showed stronger cell signalling activities with NK and B cells (Fig. 1E, Supplementary Fig. S1D, E). In summary, by employing single-cell RNA sequencing, we delineated the dynamic alterations in immune cell populations across various treatment cycles and demonstrated that when sotorasib resistance emerged, the proportions of T cells and NK cells decreased.

**Fig. 1.**
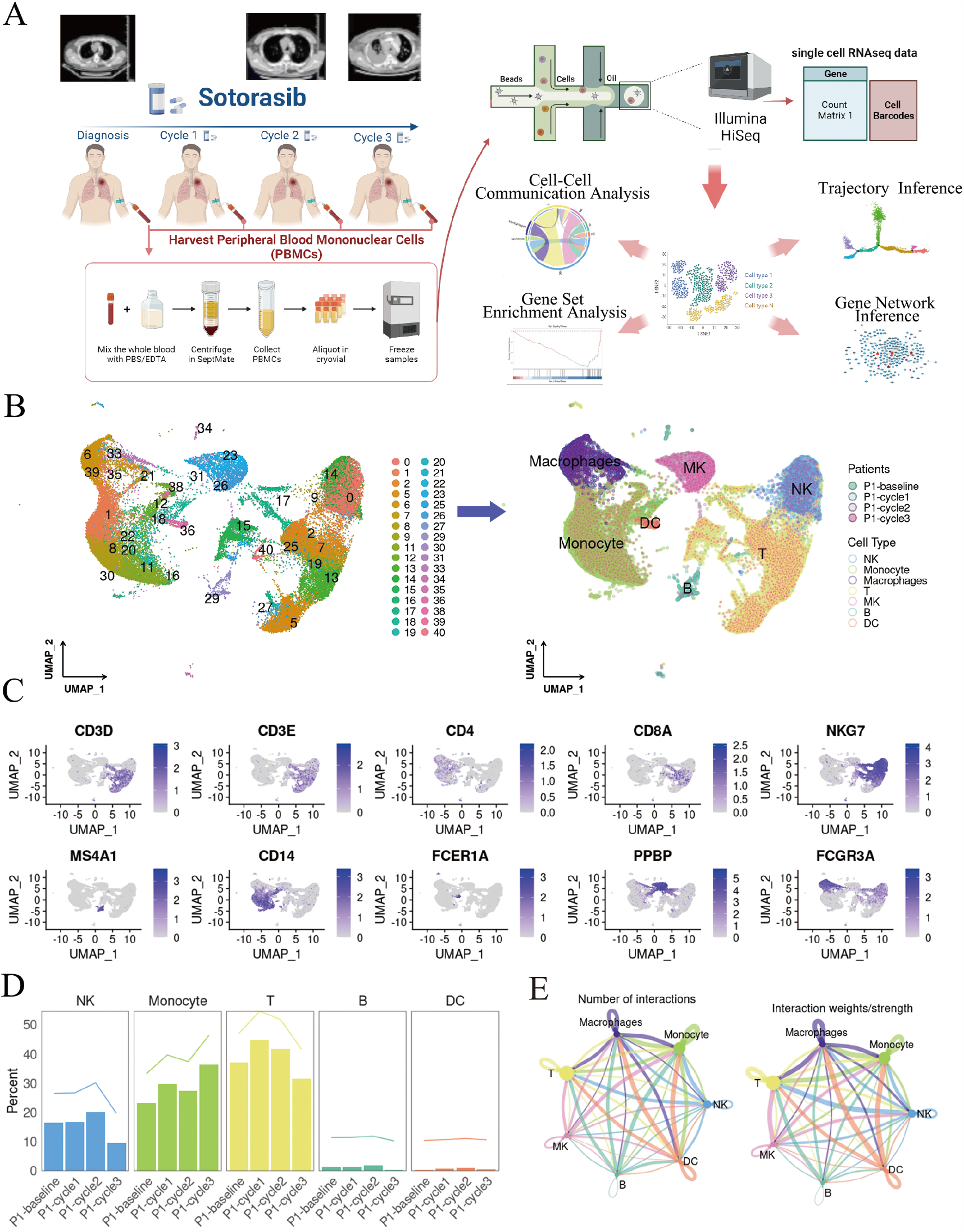
Immune cell dynamics in patient with NSCLC treated with sotorasib at different time points. (A) Schematic overview of the experimental design and analytical workflow (B) UMAP of PBMCs from the different cycles of one patient (C) Heatmap of expression levels of typical marker genes in different cells (D) Changes in the proportion of cell types in different stages (E) Communication signals activated between T cells and NK and B cells

Considering the substantial differences in T cells at different stages, we refined T cells based on expression of canonical genes, resulting in identification of 3 distinct clusters, including stem-like T cells (marked by *CD38*), CD4^+^ T cells (marked by *CD4*), and CD8^+^ T cells (marked by *CD8A*) (Supplementary Fig. S2A). Previous research has demonstrated that blood is a crucial pathway for CD8^+^ T-cell movement between secondary lymphoid organs, primary tumours, and metastases, thus offering an ideal medium for investigating peripheral antitumour responses^20,21^. Thus, we clustered CD8^+^ T cells and obtained 10 transcriptionally distinct subclusters: CD8-*FGFBP2* (Cluster 0), CD8-*TRBC1* (Cluster 1), CD8-*RPL2* (Cluster 2), CD8-*CMC1* (Cluster 3), CD8-*PPBP* (Cluster 4), CD8-*TRBV3* (Cluster 5), CD8-GNLY (Cluster 6), CD8-*GZMB* (Cluster 7), CD8-*KLRB1* (Cluster 8), and CD8-*ACTB* (Cluster 9) (Fig. 2A, B). Based on bioinformatic analysis, Clusters 2 and 4 have naive T-cell features, while others are linked to effector and cytotoxic T cells. The CD8^+^ T-cell developmental trajectory from Slingshot^22^ indicated that Clusters 2 and 4 are likely starting points, with the cells then diverging. Cluster 8 appears to have evolved from Cluster 2 directly but did not further differentiate (Fig. 2C). Analysis revealed that expression of the *KLRB1* gene, specific to Cluster 8, gradually increased during its differentiation from Cluster 2 (Supplementary Fig. S2B). Moreover, after treatment response, we observed a decrease in the percentage of the CD8-*KLRB1* cluster in the patient with KRAS^G12C^ and an increase in the percentage following treatment nonresponse (Supplementary Fig. S2C). Further investigation indicated that CD8-KLRB1 expression-stimulating cytokine genes, such as *IFNG* (IFN-γ) and *PRF1* (Perforin), are associated with cytotoxicity (Fig. 2D). In addition, the activation marker *CD69*, which is associated with mucosal-associated invariant T (MAIT) cells^12^, was found in this cluster (Fig. 2D). However, no T-cell exhaustion marker genes, such as *PDCD1* (PD-1) or *HAVCR2* (TIM-3), were detected in this cluster (Supplementary Fig. S2D). Moreover, our investigation revealed a notable expression pattern of lung-homing markers, specifically *CXCR6* and *CCR5*, within the CD8-*KLRB1* subset (Fig. 2E).

**Fig. 2.**
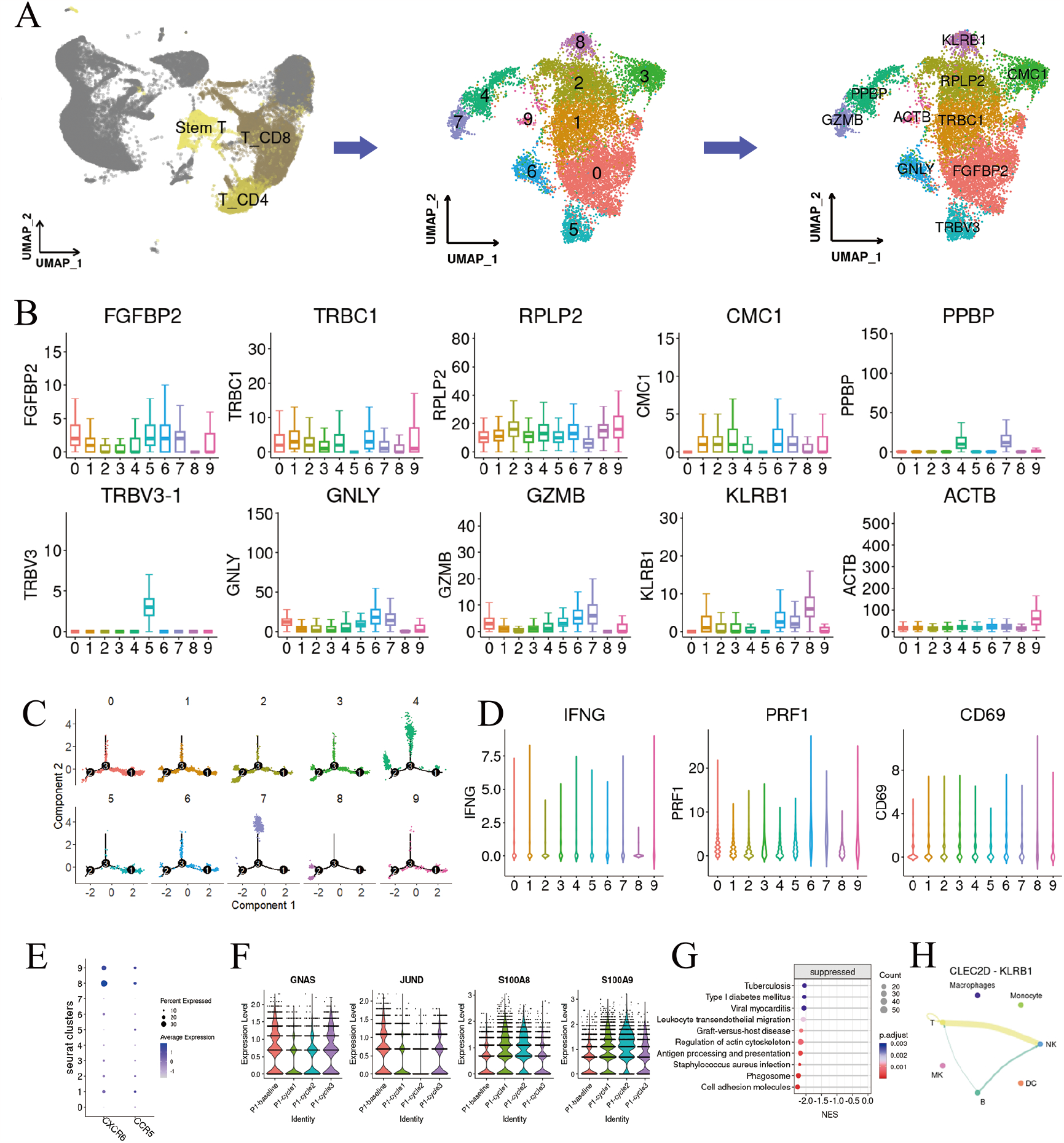
Characteristics and dynamics of CD8^+^ T-cell subsets. (A) UMAP plot after reclustering of CD8^+^ T cells in different cycles (B) Heatmap of marker genes for different subclusters of CD8^+^ T cells (C) Trajectory analysis for the CD8^+^ T-cell clusters (D) Expression of cytotoxic cytokines in CD8^+^ CD161^hi^ T cells (E) Expression of lung-homing marker genes in CD8^+^ CD161^hi^ T cells (F) Drug-related gene expression of CD8^+^ CD161^hi^ T cells (G) GSEA of CD8^+^ CD161^hi^ T cells (H) Expression levels of genes related to CLEC signaling pathway

Subsequently, within the CD8-*KLRB1* subpopulation, we identified certain genes associated with drug response. Specifically, *GNAS* and *JUND* exhibited downregulation during cycles 1 and 2, followed by upregulation in cycle 3. In contrast, *S100A8* and *S100A9* demonstrated divergent expression patterns (Fig. 2F). To investigate drug-related gene mechanisms, GSEA was used to enrich significantly changed signalling pathways among different cell clusters. The cell adhesion pathway and RAP1 pathway in the CD8-*KLRB1* cluster were suppressed (Fig. 2G). Additionally, we employed the single-cell network inference (SCENIC) approach to dissect the regulatory landscape of transcription factors within the CD8^+^ T-cell compartment^23^. Active transcription factors in CD8-*KLRB1* T cells included *JUN*, FOS, and *JUNB* (Supplementary Fig. S2E, F). GO and KEGG enrichment analyses demonstrated that the transcription factors of significance were linked to the biological processes of the MAPK signalling pathway (Supplementary Fig. S2G, H).

Previous studies have shown that lectin-like transcript 1 (*LLT1/CLEC2D*), the ligand of CD161 (encoded by *KLRB1*), is expressed on monocyte-derived dendritic cells and on activated B cells^24,25^. We detected intercellular signalling interactions between CD8-*KLRB1* T cells and natural killer (NK) and B cells mediated by the receptor-ligand pairs involving *CLEC2D* and *KLRB1* (Fig. 2H). Thus, CD8^+^ CD161^hi^ T cells and B and NK cells promote antitumour effects through the LLT1 signalling pathway.

This study presents the first comprehensive, high-resolution analysis of the peripheral blood immune landscape in patients with the KRAS^G12C^ mutation undergoing treatment with sotorasib. By leveraging single-cell technologies, we meticulously characterized the dynamic shifts within circulating PBMCs. Notably, we elucidated the intricate interplay among CD8^+^CD161^hi^ T cells and diverse immune populations, corroborating progression from naive T cells to a phenotypically distinct effector state after treatment. Moreover, our investigation revealed dynamic alterations in expression of genes associated with the pharmacological action of sotorasib. These findings provide compelling evidence of active engagement of immune effectors in the antitumour response, potentially offering novel biomarkers for monitoring treatment efficacy in real time. Tumour-infiltrating CD8^+^CD161^hi^ T cells are found in various tumour microenvironments, and their increase correlates with better prognosis^26,27^. CD161 is recognized as a coinhibitory receptor on natural killer (NK) cells; however, its role in T cells remains less defined^26,28^. Remarkably, patients with lung cancer show a higher frequency of circulating MAIT cells relative to healthy individuals^29^. Our investigation provides the first in-depth characterization of the developmental trajectories and transcriptional regulator alterations of CD8^+^CD161^hi^ T cells within the context of therapeutic intervention.

To robustly ascertain the generalizability of the observed immune response— specifically, the evolutionary dynamics of the CD8^+^CD161^hi^ T-cell subset—future investigations must include larger patient cohorts with diverse KRAS genotypes. Despite the lower prevalence of KRAS mutations in the Chinese NSCLC patient population relative to their Caucasian counterparts—with an incidence below 30%— the significant annual incidence of new cancer cases in China underscores the critical need to address this genetic aberration^30^. Thorough molecular profiling in these diverse demographic settings is essential to tailor precision oncology approaches and to enhance the clinical impact of KRAS^G12C^-targeted therapies.

## Methods

### Patient eligibility

Kiang Wu Hospital approved this study under approval number 2022-018. Blood samples and associated patient data were obtained from individuals at Macau Kiang Wu Hospital, with all participants providing their written informed consent.

### Clinical history and sample context

The patient was a 67-year-old man with stage cT1N3M1, IV (AJCC 8th) lung adenocarcinoma with the KRAS-G12C mutation. Molecular tests revealed EGFR-wildtype, PD-L1 expression was less than 1%, and ALK and Braf were negative. He refused frontline treatment chemotherapy with immunotherapy and chose therapy targeting KRAS^G12C^. Sotorasib was administered after consent was obtained. The duration of sotorasib therapy was contingent upon its effectiveness, but the mean cycle lasted approximately three to four weeks. Both regular CT scan data and clinical pathology data were recorded. The patient underwent two successive rounds of therapy. Before initiating the fourth course, an evaluation revealed tumour progression, and the patient died from the disease.

### Whole blood sample extraction and processing

Blood samples were stored in EDTA anticoagulant tubes at the clinic and processed immediately on the collection day. PBMCs and serum were isolated with standard protocols^17^. Cell viability was checked to ensure >90% viability before freezing the PBMC samples. The PBMCs were loaded onto microfluidic devices, and scRNA-seq libraries were constructed according to the Singleron GEXSCOPE protocol using GEXSCOPE Single-Cell RNA Library Kit (Singleron). Cell Ranger v 3.0 was used to generate unique molecular identifiers of genes and cellular barcodes.

### Major cell type identification and analysis

Cell type identification and clustering analysis were performed using the Seurat package (version 2.3)^31^. Cells were filtered by those that expressed fewer than 6000 genes, more than 200 genes, and less than 20% mitochondrial genes. Potential double cells were removed by DoubletFinder.

SCTransform can be used to correct the effect of sequencing depth and generate the top 2000 variant genes for principal component analysis. The first 30 main components were used with the FindClusters function, and the R package Clustree was used to visualize the evolution of cell clusters at different resolutions. We finally set the resolution to 1.1 and obtained 41 cell clusters to obtain as many cell clusters with significant differences as possible. To annotate the cell type of each cell cluster, we used SingleR to annotate each cell’s cell identity automatically. In addition, we further confirmed the cell type to which each cell cluster belongs through the following typical markers: monocytes (*CD14, LYZ, VCAN*), T cells (*CD2, CD3D, CD3E, CD3G, IL32*), NK cells (*NKG7, GNLY, KLRB1, SPON2, KLRD1, PRF1, GZMB*), macrophages (*FCGR3A*), DC cells (*CD1C, FCER1A*), B cells (*CD79A, MS4A1, CD19, CD38*), and MK cells (*PPBP, PF4*). We also noticed that a subset of cell clusters was positive for neutrophil-red blood cell markers, and these cells were removed from subsequent analyses. Comprehensive cluster analysis results of SingleR (version 2.2.0) and canonical markers, visualized by uniform prevalence approximation and projection (UMAP).

To evaluate whether the composition of primary cell types in patients differs significantly at different drug stages, we calculated their statistical significance using the scProportion (version 0.0.0.9000)^32^. We also used CellChat (version 1.6.1) to analyse and visualize interactions between major cell types. We used CellChatDB.human as a reference dataset to calculate significantly overexpressed ligands and receptors in cells to infer the biological intercellular communication network.

### Cell Subtype Identification

We clustered T cells, NK cells, monocytes, and macrophages individually. For NK cells, monocytes, and macrophages, we set the resolution to 0.4. For T cells, we used the following markers to differentiate CD4, CD8, and stem T cells: CD4 T (CD4), CD8 T (CD8A, CD8B) and stem T (CD38, CD7, PTPRC, IL7R). Since CD4 expression is lower, CD4 is depleted by CD3G+, CD3E+ and CD3D+ CD8A- and CD8B-. We reclustered CD8 T cells and obtained 10 different CD8 T subclusters at a resolution of 0.6. CellChat was used to analyse the signalling pathways activated among different cell subpopulations.

### Drug-related gene identification

Specifically, we used Presto (version 1.0.0) to identify differentially expressed genes between different medication stages. Genes with opposite expression trends between the first and third cycles were considered drug-related genes.

### Analysis of the function and regulatory mechanism of drug-related genes

We used the R package clusterProfiler (version 4.8.2)^33^ to identify the ontology terms of drug-related genes and used GSEA(version 1.62.0)^34^ to enrich the most significantly changed signaling pathways in different cell clusters. We utilized MEGENA(version 1.3.7)^35^ to construct a gene regulatory network for specific cell clusters. This was then integrated with the transcription factor regulatory network inferred by SCENIC (version 1.0) to investigate the mechanism of the drug-related genes.

### Statistical analysis

The R version was 4.3.1. A P value less than 0.05 was considered statistically significant in all analyses.

## Acknowledgement

This work was funded by regular grants (File no. 0111/2020/A3 & 0058/2020/A2) and Dr. Neher’s Biophysics Laboratory for Innovative Drug Discovery (File no. 001/2020/ALC) supported by the Macau Science and Technology Development Fund. This work was also supported by the Jointly Funded Scientific Research Project by the Ministry of Science and Technology of the People’s Republic of China, the Macao Science and Technology Development Fund (File no. 0056/2020/AMJ), the 2020 Young Qihuang Scholar funded by the National Administration of Traditional Chinese Medicine, the National Natural Science Foundation of China (82003779 and 82001995), and the FDCT Funding Scheme for Postdoctoral Researchers of Higher Education Institutions (0017/2021/APD). This work is also financially supported by the Start-up Research Grant of University of Macau (SRG2022-00020-FHS). This work is also financially supported by Multi-year research grant–general research grant of University of Macau (MYRG-GRG2023-00005-FHS-UMDF) and the Faculty of Health Science, University of Macau.

