## Supplementary figures and images for "Evolution of resistance to KRAS^G12C^ inhibitor in a non-small cell lung cancer responder"

### Supplementarydata

Supplement Fig. 1

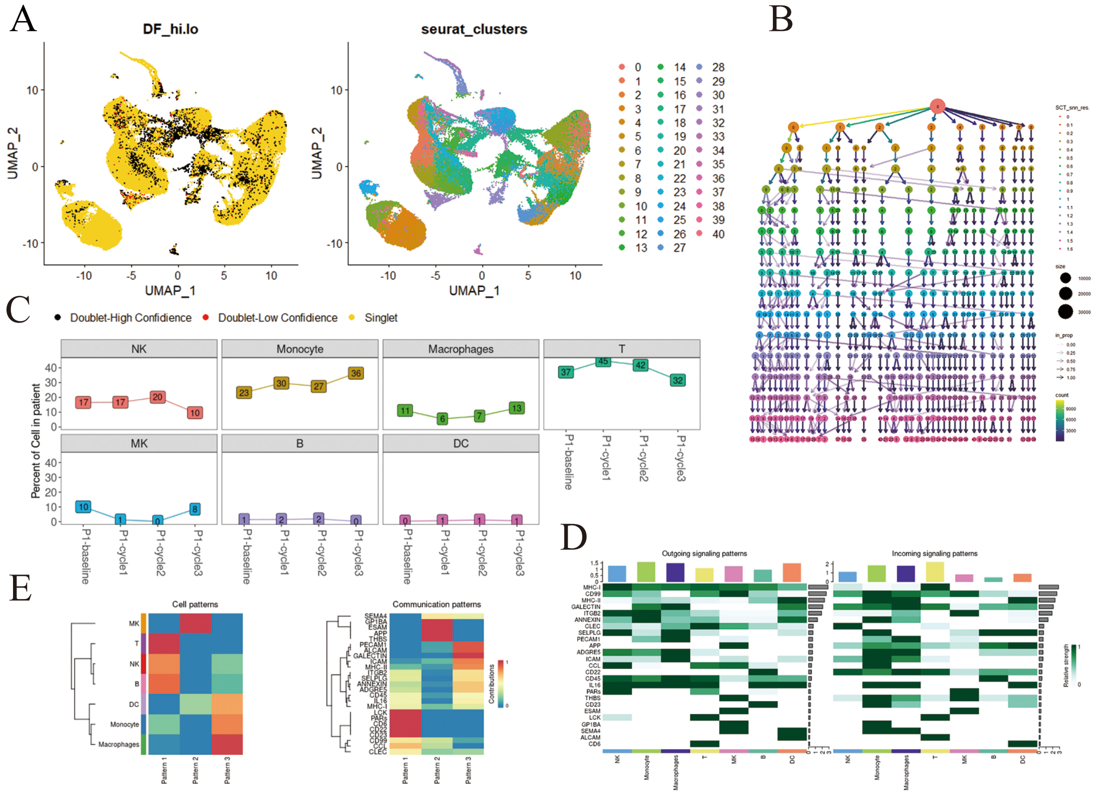


Supplement Fig. 2

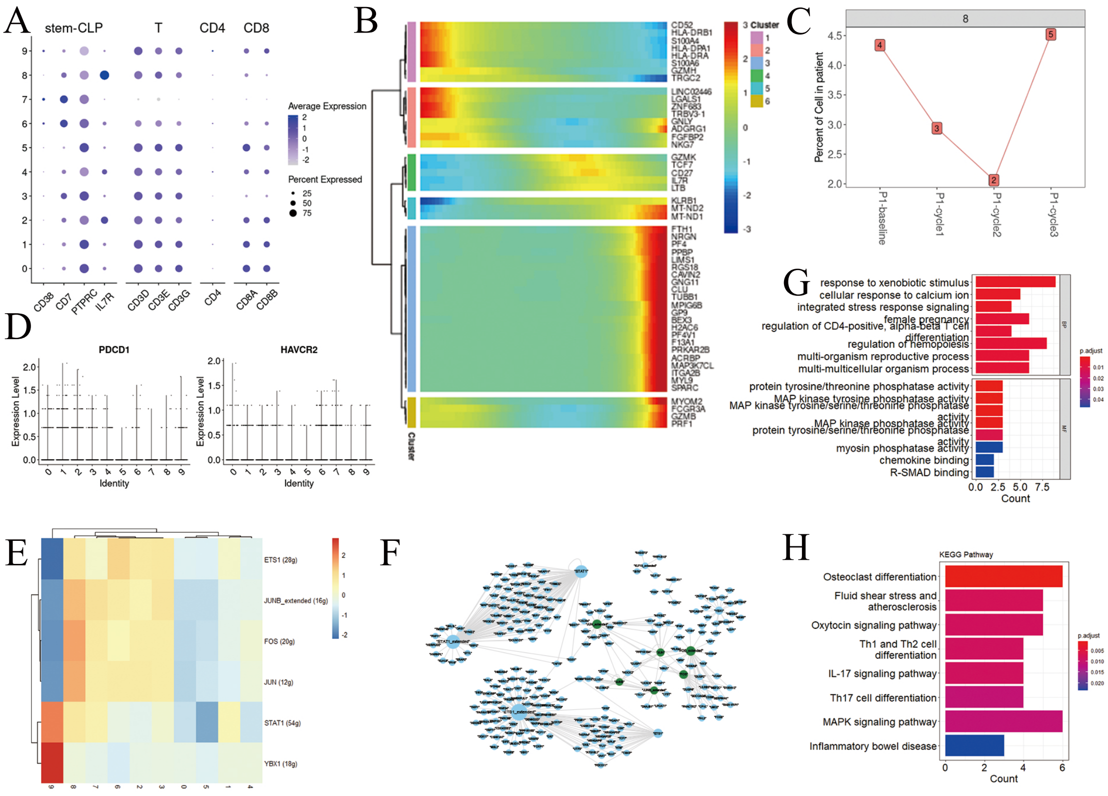
